# Religious Belief and Cognitive Conflict Sensitivity: A Preregistered fMRI Study

**DOI:** 10.1101/2020.01.16.899021

**Authors:** Suzanne Hoogeveen, Lukas Snoek, Michiel van Elk

**Affiliations:** Department of Psychology, University of Amsterdam, The Netherlands

**Author notes:** Correspondence concerning this article should be addressed to: Suzanne Hoogeveen, Nieuwe Achtergracht 129B, 1001 NK Amsterdam, The Netherlands.

**Keywords:** Religiosity, Cognitive conflict, Functional magnetic resonance imaging, Anterior cingulate cortex

## Abstract

In the current preregistered fMRI study, we investigated the relationship between religiosity and behavioral and neural mechanisms of conflict processing, as a conceptual replication of the study by Inzlicht et al. (2009). Participants (*N* = 193) performed a gender-Stroop task and afterwards completed standardized measures to assess their religiosity. As expected, the task induced cognitive conflict at the behavioral level and at a neural level this was reflected in increased activity in the anterior cingulate cortex (ACC). However, individual differences in religiosity were not related to performance on the Stroop task as measured in accuracy and interference effects, nor to neural markers of response conflict (correct responses vs. errors) or informational conflict (congruent vs. incongruent stimuli). Overall, we obtained moderate to strong evidence in favor of the null hypotheses that religiosity is unrelated to cognitive conflict sensitivity. We discuss the implications for the neuroscience of religion and emphasize the importance of designing studies that more directly implicate religious concepts and behaviors in an ecologically valid manner.

---

Everywhere across the world, in all times and cultures we find people who believe in super-natural beings. Religious beliefs seem highly successful in offering explanations for various phenomena, ranging from how the world originated, to why one had to switch jobs and what happens after one dies. Yet these beliefs are difficult - if not impossible - to support with empirical evidence. In fact, believers are often confronted with widely supported contradicting evidence, for instance evolutionary explanations of the origins of life or reductionistic explanations of their religious experiences. And yet, despite these challenges, most religious believers keep up their faith (Pew Research Center, 2012).

Various scholars have suggested that a mechanism of reduced conflict sensitivity, i.e., detecting the incongruency between two potentially conflicting sources of information, may foster the acceptance and maintenance of religious beliefs. For example, dual-process accounts of religion (Risen, 2016), the predictive processing model (van Elk & Aleman, 2017), and the cognitive resource depletion model (Schjoedt et al., 2013) all assume that religiosity is associated with a reduced tendency for analytical thinking and error monitoring. A process of reduced conflict detection (or correction) makes individuals less prone to note information that seemingly contradicts their religious worldviews and to update their beliefs in the light of new information. This tendency could potentially underlie the relative immunity of religious beliefs to criticism based on empirical observations (cf. what van Leeuwen, 2014 calls ‘evidential invulnerability’).

Notably, the implicit assumption of most theoretical frameworks appears to be that a mechanism of reduced conflict sensitivity makes people more receptive to being religious. However, it could also be that being religious affects people’s sensitivity to conflicting information; religious ‘training’ inoculates believers against contractions and violations of their worldview. This notion parallels findings from mindfulness meditation research reporting evidence that meditation training increases cognitive control as it teaches practitioners to suppress irrelevant information (Moore & Malinowski, 2009; Teper & Inzlicht, 2012), with meditation experts showing less activation in brain areas implicated in attention and cognitive control (e.g., the anterior cingulate cortex; Brefczynski-Lewis, Lutz, Schaefer, Levinson, & Davidson, 2007). However, whereas mindfulness meditation may train practitioners to flexibly suppress irrelevant information –resulting in increased cognitive control– religious training may particularly enhance the salience of intuitive information, while suppressing contradicting analytical alternatives.

In line with this suggestion, several empirical studies found that increased religiosity is related to a decreased cognitive performance, especially when a logically correct response must override a conflicting intuitive response (e.g., in a base-rate fallacy test; Daws & Hampshire, 2017; Good, Inzlicht, & Larson, 2015; Pennycook, Cheyne, Barr, Koehler, & Fugelsang, 2014; Zmigrod, Rentfrow, Zmigrod, & Robbins, 2018). Other behavioral studies correlated individuals’ self-reported level of religiosity with their performance on low-level cognitive control tasks such as the Go/No-go task or the Stroop task. These studies present a mixed bag of evidence; some report a positive relationship (Inzlicht, McGregor, Hirsh, & Nash, 2009), an inconsistent pattern (Inzlicht & Tullett, 2010), or no relation-ship (Kossowska, Szwed, Wronka, Czarnek, & Wyczesany, 2016) between religiosity and cognitive control (in terms of accuracy and reaction times).

In addition to this behavioral research, a few neuroscientific studies have been conducted on the association between religiosity and conflict sensitivity. For instance, an fMRI study investigated brain responses in devoted religious believers who listened to intercessory prayer. When participants believed that the prayer was pronounced by a charismatic religious authority, they showed a reduced activation of their frontal executive network, including the dorsolateral prefrontal cortex (DLPC) and the ACC, which have been associated with conflict detection (Schjoedt, Stødkilde-Jørgensen, Geertz, Lund, & Roepstorff, 2011). Furthermore, Inzlicht et al. (2009) conducted a series of EEG studies looking at the relation between religiosity and the error-related negativity (ERN; Inzlicht et al., 2009; Inzlicht & Tullett, 2010). Compared to skeptics, religious believers demonstrated a smaller ERN amplitude in response to errors on a color-word Stroop task (Inzlicht et al., 2009). The authors suggest that these findings reflect the palliative effects of religiosity on distress responses: religious believers experience less distress in association with committing an error and this is reflected in a reduced ERN amplitude. There is, however, an open-ended debate on the functional significance of the ERN; while some researchers interpret the ERN primarily as an affective (i.e., distress) signal, others emphasize that it mainly reflects conflict-sensitivity (Yeung, Botvinick, & Cohen, 2004; Bush, Luu, & Posner, 2000; Hajcak, Moser, Yeung, & Simons, 2005; Botvinick, Braver, Barch, Carter, & Cohen, 2001; Maier & Steinhauser, 2016; Carter et al., 1998).

Relatedly, different views have been proposed on how the relation between religiosity and ACC conflict activity should be interpreted; whereas Inzlicht, Tullett, and Good (2011) suggest that ACC activity in this context reflects error distress, Schjoedt and Bulbulia (2011) argue that the interpretation of ACC activity as reflecting purely cognitive conflict sensitivity is more parsimonious. We believe this discussion partly hinges upon the operationalisation of ‘conflict’. EEG studies on cognitive conflict have typically studied the ERN as a proxy for ACC activity. The ERN is an *error*-related signal and reflects neural activity associated with incorrect vs. correct responses, i.e., conflict at the level of the behavioral response (hereafter: response conflict). In contrast, fMRI studies on cognitive conflict typically focus on the neural activity associated with incongruent vs. congruent stimulus trials, i.e., conflict at the level of information processing (hereafter: informational conflict). Although there is often a correlation between response conflict^1^ and informational conflict, not all incongruent trials result in errors, nor do all congruent trials by definition result in correct responses. It is therefore important to dissociate between these two levels of conflict and their associated neural activity (cf. Tang, Critchley, Glaser, Dolan, & Butterworth, 2006; van Veen & Carter, 2005).

It thus remains unclear to what extent religiosity is related to a reduced sensitivity for response conflict (e.g., responding with ‘green’ when it should have been ‘red’) or to a reduced sensitivity for informational conflict (e.g., seeing the word ‘green’ printed in a red font). An effect for *response conflict* should be reflected in a relationship between religiosity and the strength of the error–correct Stroop contrast in the fMRI data, which would be a direct replication of the study by Inzlicht et al. (2009) and their proposed framework (Inzlicht et al., 2011; Proulx, Inzlicht, & Harmon-Jones, 2012). An effect for *informational conflict* should be reflected in a relationship between religiosity and the strength of the incongruent–congruent Stroop contrast in the fMRI data. Schjoedt and Bulbulia (2011), for instance, indeed seem to interpret Inzlicht et al.’s results as religious believers’ inattention to conflict monitoring. In everyday life, both sources of conflict detection could play a role in the maintenance of religious beliefs, e.g., when a believer simply does not detect the incongruency between different sources of information or when he / she fails to suppress an intuitive but objectively incorrect answer.

Taking the distinction between response conflict and informational conflict into account, here we investigated two different hypotheses regarding the relation between religiosity and cognitive conflict sensitivity: (1) there is a negative relationship between religiosity and ACC activity induced by response conflict (i.e., the incorrect–correct response contrast), and (2) there is a negative relationship between religiosity and ACC activity induced by informational conflict (i.e., the incongruent–congruent Stroop contrast). We note that both hypotheses are not mutually exclusive, as religiosity could be related to both mechanisms of conflict detection.^2^

Although earlier studies provide preliminary evidence for the religiosity–conflict sensitivity relation, we believe the present study –including a conceptual replication of the seminal study by Inzlicht et al. (2009)– is important for the following reasons. First, in order to substantiate the notion that religious believers are characterized by a *general* tendency for reduced conflict sensitivity at the neural level, a significant correlation or inter-group difference should be established. So far, only two studies found evidence for an inverse relation between religiosity and conflict-induced ACC activity (Inzlicht et al., 2009; Kossowska et al., 2016), while one study failed to find such a relation (Good et al., 2015). Second, with the exception of Good et al. (2015, *n* = 108), all experiments linking religiosity to ACC activity included small samples and were therefore most likely underpowered (i.e., Inzlicht et al., 2009, *n* = 28 [Study 1], *n* = 22 [Study 2]; Kossowska et al., 2016, *n* = 37) Third, the hypothesized relation between religiosity and cognitive conflict is primarily based on either behavioral or EEG data. EEG studies, however, can offer only indirect evidence for the involvement of specific brain areas (Gazzaniga & Ivry, 2013). The use of fMRI may complement the existing findings, as fMRI allows for a higher spatial specificity, and may thus provide more conclusive evidence regarding the role of the ACC in the acceptance and maintenance of religious beliefs. Finally, the current study design allowed us to dissociate between neural effects related to response conflict (i.e., activity predicted by response accuracy) and to informational conflict (i.e., activity predicted by Stroop congruency). This may help to disentangle the ‘conflict sensitivity’ accounts of religiosity, and hence affords a more precise theoretical interpretation of the existing data.

## Hypotheses

We tested eight hypotheses, four of which were based on our research questions and four that served as ‘outcome neutral tests’ (Chambers, Feredoes, Muthukumaraswamy, & Etchells, 2014). The four outcome neutral tests were used to validate that our task did indeed induce cognitive conflict (reflected in accuracy and Stroop interference effects), that error commission was reflected in ACC activity, and that informational conflict was reflected in ACC activity. The corresponding outcome neutral hypotheses for the behavioral measures were: (ℋ_1_) participants are more accurate on congruent compared to incongruent Stroop trials, and (ℋ_2_) participants respond faster on congruent compared to incongruent Stroop trials. Outcome neutral hypotheses for the neural measures were: (ℋ_3_) errors on the Stroop task induce more ACC activity compared to correct responses, on average across subjects, and (ℋ_4_) incongruent Stroop trials induce more ACC activity compared to congruent trials, on average across subjects.

Conditional on establishing the effects related to hypotheses 1–4, we tested four corresponding hypotheses about the relation between religiosity and conflict sensitivity. For the behavioral measures, we hypothesized that (ℋ_5_) Stroop accuracy is negatively related to religiosity, and (ℋ_6_) Stroop interference (i.e., the difference in RT for incongruent vs. congruent trials) is positively related to religiosity, indicating decreased cognitive performance. We note that, based on the existing literature one could hypothesize both a positive and a negative relationship between religiosity and conflict detection; on the one hand, religiosity is associated with reduced response conflict and hence smaller interference effects (cf. Inzlicht et al., 2011). On the other hand, religiosity is associated with an increased tendency for intuitive responding, which means that more effort is required to overcome these intuitive response on incongruent Stroop trials, hence larger interference effects should be expected (cf. Pennycook et al., 2014). Despite these divergent theoretical predictions, most studies have not found any association between religiosity and Stroop interference (Inzlicht et al., 2009, Study 1; Inzlicht & Tullett, 2010; Kossowska et al., 2016), except for Study 2 by Inzlicht et al. (2009), in which a positive correlation between religiosity and Stroop interference was reported. Here, in line with the latter finding we hypothesized a positive relationship between religiosity and Stroop interference.

For the neural measures, we hypothesized that (ℋ_7_) the size of the error–correct response BOLD signal contrast (i.e., difference in BOLD signal between errors and correct responses) in the ACC is negatively related to religiosity, on average across subjects (cf. Inzlicht et al., 2009), and (ℋ_8_) the size of the incongruent–congruent BOLD signal contrast (i.e., difference in BOLD signal between the incongruent and congruent condition) in the ACC is negatively related to religiosity, on average across subjects. All hypotheses were pre-registered on the Open Science Framework (see https://osf.io/nspxb/registrations). Finally, we added exploratory whole-brain analyses to explore whether religiosity is associated with conflict-induced neural activity in any other brain areas besides the ACC.

## Methods

### Overview

The data for this study had already been collected as part of the Population Imaging (PIoP) project (May 2015 - April 2016), conducted at the Spinoza Center for Neuroimaging at the University of Amsterdam (see Appendix A for a description of the project). An overview of the data collection and analysis procedure is presented in Figure 1. All hypotheses were formulated independently without any knowledge of the preprocessed data, and the analysis pipeline was developed and preregistered prior to data inspection.^3^ The preregistration can be accessed on the OSF (https://osf.io/nspxb/). This folder also contains the anonymized raw and processed data and the R scripts used to preprocess the behavioral data and to conduct the confirmatory analyses (including all figures). The preprocessing scripts for the fMRI analysis and the exploratory fMRI analyses can be found at https://github.com/lukassnoek/ReligiosityFMRI. The (uncorrected) brain maps can be found at https://neurovault.org/collections/6139/.

**Figure 1.**
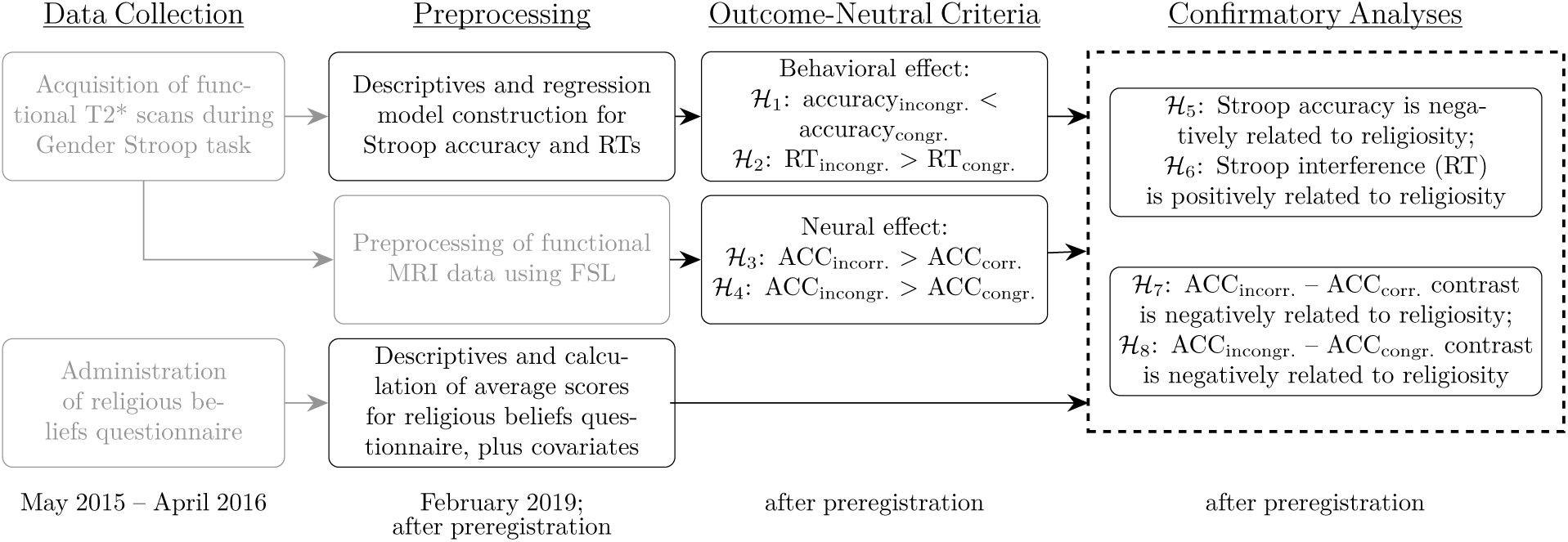
Overview of data acquisition and analysis. Boxes marked in grey had already been completed prior to commencing this project. Boxed marked in black represent the analysis steps for the present study, which were determined in the preregistation.

### Participants

Participants were students who were recruited at the University of Amsterdam and received a financial remuneration. Participants were screened for MRI contraindications before MRI data acquisition. The intended number of participants was 250, but due to technical problems during part of the acquisition process, only 244 participants yielded usable MRI data. Of those 244, data from 20 subjects were excluded due to artifacts in the MRI data due to scanner instabilities or errors during export and/or reconstruction of the data. Additionally, 10 participants were excluded because they did not complete the task of interest (i.e., the gender-Stroop task). These exclusions were known at the time of the preregistration.

We entered the analysis phase with data from *N* = 214 participants. Out of these 214, eight participants were excluded –as preregistered– because they did not complete the religiosity questionnaire or lacked data on the covariates of interest (age, gender, and intelligence). We additionally preregistered to exclude participants whose accuracy was lower than 65%, because this indicates performance at chance level. This means that participants who responded correctly on fewer than 63 out of the 96 trials were excluded. Furthermore, participants who did not respond within the response interval on more than 20% of the Stroop trials were also excluded. As the minimum response interval of 4500ms is assumed to be sufficient for timely responses, missed responses on more than 20% of the trials were taken to indicate that participants did not understand or perform the task adequately. These criteria led to the exclusion of 14 participants, yielding a total sample size of 193. In addition, for the fMRI analyses, there were 21 participants who did not make any mistake during the task, preventing us from calculating the ‘incorrect–correct’ contrast.^4^ As such, the confirmatory ROI and whole-brain analyses of this contrast were based on data from 172 participants. All other analyses were done on a total of *N* = 193 participants with complete data. The final sample consisted of 109 (56.5%) women and 84 (43.5%) men. The average age of the participants was 22.2 years (*SD* = 1.9; range = 18 − 26).

The study was approved by the local ethics committee at the Psychology Department of the University of Amsterdam (Project #2015-EXT-4366) and all participants were treated in accordance with the Declaration of Helsinki.

#### Sample Size Justification

The sample size was determined based on the target of the overall project minus exclusions due to artifacts in the data, incomplete data, or preregistered quality criteria. As there were no existing fMRI studies on the relation between religiosity and cognitive conflict processing –only EEG studies– we could not perform a power analysis. However, we note that a sample of *N* ≈ 200 is substantially large for an fMRI study (Szucs & Ioannidis, 2017)^5^ and exceeds the recommended minimum sample size of *N* = 100 for correlational (neuroimaging) research (Dubois & Adolphs, 2016; Schönbrodt & Perugini, 2013).

### Procedure

The study ran from May 2015 until April 2016. On each testing day, two participants were tested, which took approximately 4 hours and included an extensive behavioral test battery (approximately 2.5 hours) and an MRI session (approximately 1.5 hours). The order of behavioral and MRI sessions were counterbalanced across participants.

### Study Design

The study involved a mixed design with Stroop congruency as the within-subjects variable and religiosity as the between-subjects continuous individual differences variable. The main part of the study qualified as an observational study; we investigated the correlation between performance on the Stroop task and religiosity, and between BOLD-fMRI activity and religiosity, without manipulating any variables except for trial congruency (congruent vs. incongruent Stroop trials). The fMRI task involved a rapid event-related design; a hypothesized BOLD response was modelled following the presentation of facial stimuli in the congruent or incongruent condition, as well as following correct and incorrect responses.

### Stroop Task

We used a face-gender variant of the Stroop task (adapted from Egner, Ely, & Grinband, 2010), often referred to as the ‘gender-Stroop’ task, in which pictures of faces from either gender are paired with the corresponding (i.e., congruent) or opposite (i.e., incongruent) gender label (see below for details on the task and example pictures of the stimuli). The face-gender variant of the Stroop task (Egner & Hirsch, 2005) has been shown to induce significant behavioral conflict and neural ACC activation (Egner, Etkin, Gale, & Hirsch, 2008).^6^ Each trial consisted of a photographic stimulus depicting either a male or female face, with the gender label ‘MAN’ or ‘WOMAN’ superimposed in red, resulting in gender-congruent and gender-incongruent stimuli The Stroop condition –congruent vs. incongruent– thus formed the within-subjects manipulated variable.

The stimuli set consisted of a total of 12 female and 12 male faces, with the labels ‘man’, ‘sir’,’woman’, and ‘lady’, both in lower- and uppercase added to the pictures (e.g., ‘sir’ and ‘SIR’).^7^ All combinations appeared exactly one time, resulting in 96 unique trials (48 congruent and 48 incongruent). Participants were always instructed to respond to the gender of the pictured face, ignoring the distractor word.

The stimuli were presented for 500ms with a variable inter-trial interval ranging between 4000-6000ms, in steps of 500ms. Participants could respond from the beginning of the stimulus presentation until the end of the inter-trial interval (i.e., minimum response interval was 4500, maximum response interval was 6500), using their left and right index finger. If no button was pressed during this interval, the trial was recorded as a ‘miss’. Stimuli were presented using Presentation (Neurobehavioral Systems, www.neurobs.com), and displayed on a back-projection screen that was viewed by the subjects via a mirror attached to the head coil.

### Religiosity Measures

Our religiosity measure consisted of 7 items that were based on religiosity questions included in the World Values Survey (WVS; World Values Survey, 2010), covering religious identification, beliefs, values, and behaviors (institutionalized such as church attendance and private such as prayer). Besides having high face-validity, these measures have been validated in other studies (Lindeman, Svedholm-Hakkinen, & Lipsanen, 2015; Norenzayan, Gervais, & Trzesniewski, 2012; Stavrova, 2015) and the items have been used in previous studies (Maij et al., 2017; van Elk & Snoek, 2019). The items were evaluated on a 5-point Likert scale ranging from 1 = *not at all* to 5 = *very much*; see Table 1 for the exact items. Ratings on the seven religiosity items were tallied to create an average religiosity score per participant. For the analyses, these average scores were standardized. As anticipated in the preregistration, the distribution of the religiosity data was indeed positively-skewed, since our sample consisted of highly-secularized students. Although non-normality may reduce statistical power (Poldrack, Mumford, & Nichols, 2011), it does not pose a problem for our analysis, since Bayesian linear regression models –like general(ized) linear models in general– do not assume normality of predictors (solely of model residuals).

**Table 1.**
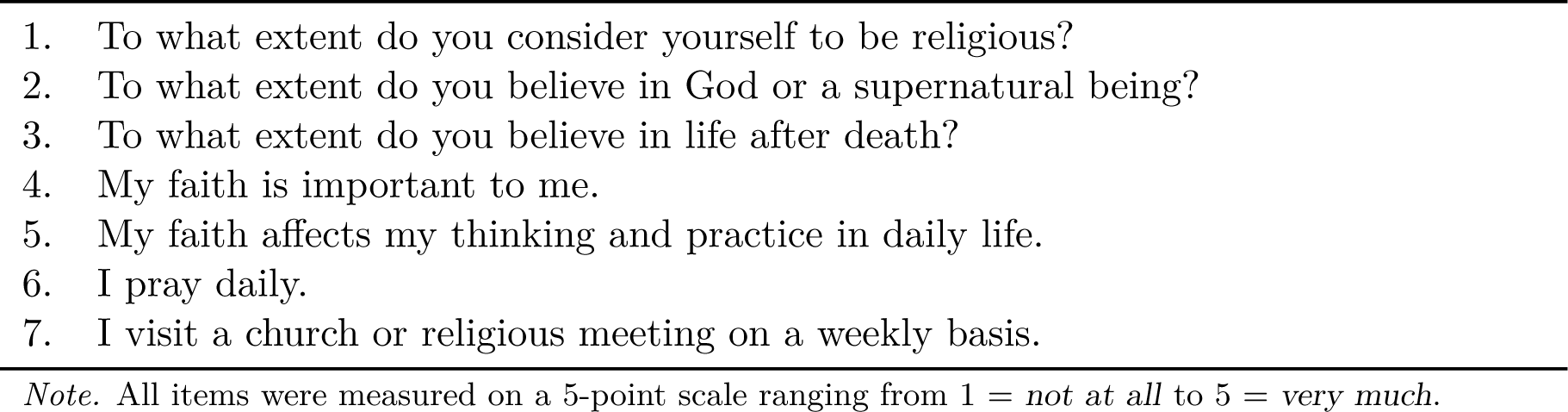
Items of the religiosity scale.

### Additional Variables

Gender, age, and intelligence were included as covariates in the analyses of the main hypotheses. Intelligence was indexed by the sum score on the (short-form) Raven matrices test (Raven, 2000). The rationale for including these measures as covariates in our analysis was to control for the potential confound that any religiosity effect may be driven by other individual differences that are known to be associated with religiosity; females are typically more religious than males (Miller & Hoffmann, 1995), older people tend to be more religious than younger people (Argue, Johnson, & White, 1999), and people scoring high on intelligence are on average less religious (Zuckerman, Silberman, & Hall, 2013). Age and intelligence scores were standardized in the analyses.

Since the proposed study was part of a larger project, a number of extra tasks and questionnaires were administered to the participants (see Appendix A for a description). These measures were not included in the present study.

### fMRI Data Acquisition

Subjects were tested using a Philips Achieva 3T MRI scanner and a 32-channel SENSE headcoil. A survey scan was made for spatial planning of the subsequent scans. After the survey scan, five functional (T2*-weighted BOLD-fMRI) scans (corresponding to five different tasks, including the gender-Stroop task; see Appendix A for an overview of the other tasks), one structural (T1-weighted) scan, and one diffusion-weighted (DWI) scan were acquired. The DWI scan will not be described further, as it is not relevant to the current study. The Stroop task was done during the second scan of the session (not including the survey scan).

The structural T1-weighted scan was acquired using 3D fast field echo (TR: 82 ms, TE: 38 ms, flip angle: 8°, FOV: 240 × 18 mm, 220 slices acquired using single-shot ascending slice order and a voxel size of 1.0 × 1.0 × 1.0 mm). The functional T2*-weighted gradient echo sequences (single shot, echo planar imaging) were run. The following parameters were used for the MRI sequence during the gender-Stroop task: TR=2000 ms, TE=27.63 ms, flip angle: 76.1°, FOV: 240 × 240 mm, in-plane resolution 64 × 64, 37 slices (with ascending slice acquisition), slice thickness 3 mm, slice gap 0.3 mm, voxel size 3 × 3 × 3 mm), covering the entire brain. During the Stroop task, 245 volumes were acquired.

### Preprocessing

Preprocessing was performed using `fmriprep` version 1.0.15 (Esteban et al., 2019, 2018), a Nipype (Gorgolewski et al., 2011, 2017) based tool. `fmriprep` was run using the package’s Docker interface. Each T1w (T1-weighted) volume was corrected for INU (intensity non-uniformity) using `N4BiasFieldCorrection` v2.1.0 (Tustison et al., 2010) and skull-stripped using `antsBrainExtraction.sh` v2.1.0 (using the OASIS template). Brain surfaces were reconstructed using recon-all from FreeSurfer v6.0.1 (Dale, Fischl, & Sereno, 1999), and the brain mask estimated previously was refined with a custom variation of the method to reconcile ANTs-derived and FreeSurfer-derived segmentations of the cortical gray-matter of Mindboggle (Klein et al., 2017). Spatial normalization to the ICBM 152 Nonlinear Asymmetrical template version 2009c (Fonov, Evans, McKinstry, Almli, & Collins, 2009) was performed through nonlinear registration with the `antsRegistration` tool of ANTs v2.1.0 (Avants, Epstein, Grossman, & Gee, 2008), using brain-extracted versions of both T1w volume and template. Brain tissue segmentation of cerebrospinal fluid (CSF), white-matter (WM) and gray-matter (GM) was performed on the brain-extracted T1w using fast (Zhang, Brady, & Smith, 2001; FSL v5.0.9).

Functional data was motion corrected using `mcflirt` (Jenkinson, Bannister, Brady, & Smith, 2002; FSL v5.0.9). ‘Fieldmap-less’ distortion correction was performed by co-registering the functional image to the same-subject T1w image with intensity inverted (Wang et al., 2017; Huntenburg, 2014) constrained with an average fieldmap template (Treiber et al., 2016), implemented with `antsRegistration` (ANTs). This was followed by co-registration to the corresponding T1w using boundary-based registration (Greve & Fischl, 2009) with 9 degrees of freedom, using `bbregister` (FreeSurfer v6.0.1). Motion correcting transformations, field distortion correcting warp, BOLD-to-T1w transformation and T1w-to-template (MNI) warp were concatenated and applied in a single step using `antsApplyTransforms` (ANTs v2.1.0) using Lanczos interpolation. Functional data was smoothed with a 5 mm FWHM Gaussian kernel. Many internal operations of `fmriprep` use Nilearn (Abraham et al., 2014), principally within the BOLD-processing workflow. For more details of the pipeline see http://fmriprep.readthedocs.io.

#### Quality Control

After preprocessing, the `MRIQC` package (Esteban et al., 2017) was used to generate visual reports of the data and results of several intermediate preprocessing steps. These reports were visually checked for image artifacts, such as ghosting, excessive motion, and reconstruction errors. Participants displaying such issues were excluded from further analysis.

#### fMRI First-Level Model

The fMRI timeseries were modelled using a first level (i.e., subject-specific) GLM, using the implementation provided by the `nistats` Python package (https://nistats.github.io; Abraham et al., 2014; version rel0.0.1b). The GLM included four predictors modelling elements of the task: incongruent trials, congruent trials, correct trials, and incorrect trials. If a participant did not make any mistakes, the ‘incorrect trials’ predictor was left out. The predictors were convolved with a canonical hemodynamic response function (HRF; Glover, 1999). Onsets for the (in)congruent trial predictors were defined at the onset of the image and had a fixed duration of 0.5 seconds. Onsets for the (in)correct trial predictors were defined at the onset of the response. Additionally, six motion regressors (reflecting the translation and rotation parameters in three dimensions) were included as covariates. GLMs were fit with AR1 autocorrelation correction. After fitting the GLMs, the following contrasts were computed: ‘incorrect–correct’ and ‘incongruent– congruent’. The parameters –beta parameters– and associated variance terms from these contrasts were used in subsequent confirmatory ROI analyses and exploratory whole-brain analyses.^8^

#### fMRI Group-Level Model (exploratory)

In addition to the confirmatory analyses, we also performed an exploratory whole-brain analysis of the effect of religiosity on fMRI activity associated with response conflict (i.e,. ℋ_7_) and informational conflict (i.e., ℋ_8_). Similar to the confirmatory analyses, in addition to religiosity, the variables age, gender, and intelligence were added as covariates to the model. In the group-level model and in accordance with the ‘summary statistics approach’, the first-level ‘incorrect–correct’ and ‘incongruent–congruent’ contrast estimates represent the dependent variables, while religiosity, age, gender, and intelligence represent the independent variables. For the participants who did not make any error, we could not compute the ‘incorrect-correct’ contrast and they were thus excluded from the group-analysis of the ‘incorrect-correct’ contrast.

We used the FSL tool `randomise` (Winkler, Ridgway, Webster, Smith, & Nichols, 2014) in combination with `threshold-free cluster enhancement` (Smith & Nichols, 2009) to perform a non-parametric group-analysis of the effect of religiosity. We ran 10, 000 permutations. Specifically, we tested for a non-directional (two-tailed) effect of religiosity variable (controlled for the other covariates). In addition, as ‘outcome neutral tests’, we computed the average of the first-level contrasts (‘intercept-only’ model) for both the ‘incorrect-correct’ and ‘incongruent-congruent’ first-level contrasts. We corrected for multiple comparisons using the distribution of the ‘maximum statistic’ under the null-hypothesis (i.e., the default in `randomise`) with a voxel-level *α* value of 0.025 (i.e., *α* = 0.05 but corrected for two-sided tests; Chen et al., 2018). We plotted the significant voxels showing either a negative or positive effect of religiosity on a standard MNI152 brain.

### ROI Definition

For this study’s confirmatory ROI analyses, we used a preregistered ROI based on a conjunction of a functional ROI, derived from fMRI activity preferentially associated with ‘error’ (for ℋ_3_ and ℋ_7_) or ‘conflict’ (for ℋ_4_ and ℋ_8_) extracted using Neurosynth (Yarkoni, Poldrack, Nichols, Van Essen, & Wager, 2011), and an anatomical ROI based on the anatomical coordinates of the ACC, taken from the Harvard-Oxford cortical atlas (Craddock, James, Holtzheimer, Hu, & Mayberg, 2012). The reasons for using a mask based on both a functional and anatomical ROI are twofold. First, the anatomical ROI of the ACC in the Harvard-Oxford atlas (and many others) consists of several putatively functionally different subregions (Vogt, 2005; Holroyd et al., 2004; Gasquoine, 2013). A functional ROI based on the Neurosynth database would resolve this issue of functional ambiguity within a single (anatomical) ROI; however, the Neurosynth maps for ‘error’ and ‘conflict’ contain more brain areas than just the ACC (such as the bilateral insula). Therefore, by using the conjunction between the functional ROIs based on Neurosynth and the anatomical ROI of the ACC, we restrict our analyses to a single *anatomical* region that is most likely to be *functionally* relevant for the psychological constructs of interest, i.e., response conflict (“error”) and informational conflict (“conflict”). We realize that due to the ambiguity of the term ‘conflict’ (which may refer to informational conflict or response conflict), the Neurosynth map for ‘conflict’ will likely also be based on studies involving response conflict. Although not ideal, we believe that this method is the most appropriate way to define our ROI.

Specifically, for our functional ROI, we used the Neurosynth Python package to conduct separate meta-analyses of the terms “error*” and “conflict*”, with a frequency threshold of 0.001^9^. We used the ‘association test map’ from the meta-analysis output (FDR-thresholded for multiple comparisons at *p <* 0.01), which reflects voxels which are *preferentially* associated with the term ‘error’ and ‘conflict’, rather than other psychological constructs. For our anatomical ROI, we used the ‘anterior cingulate cortex’ region within the Harvard-Oxford cortical atlas. We will define the ACC within this probabilistic atlas as the set of voxels with a nonzero probability of belonging to the ACC. Our final ROI is based on the logical conjunction of these two ROIs (see Figure 2). For the confirmatory ROI analyses, we averaged the GLM parameters (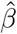, ‘beta-values’) and associated variance parameters 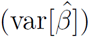 separately for the ‘incorrect–correct’ (ℋ_3_ and ℋ_7_) and ‘incongruent– congruent’ (for ℋ_4_ and ℋ_8_) first-level contrasts for each participant. These ROI-average parameters were subsequently analyzed in a hierarchical Bayesian regression model (see Statistical Models section for details).

**Figure 2.**
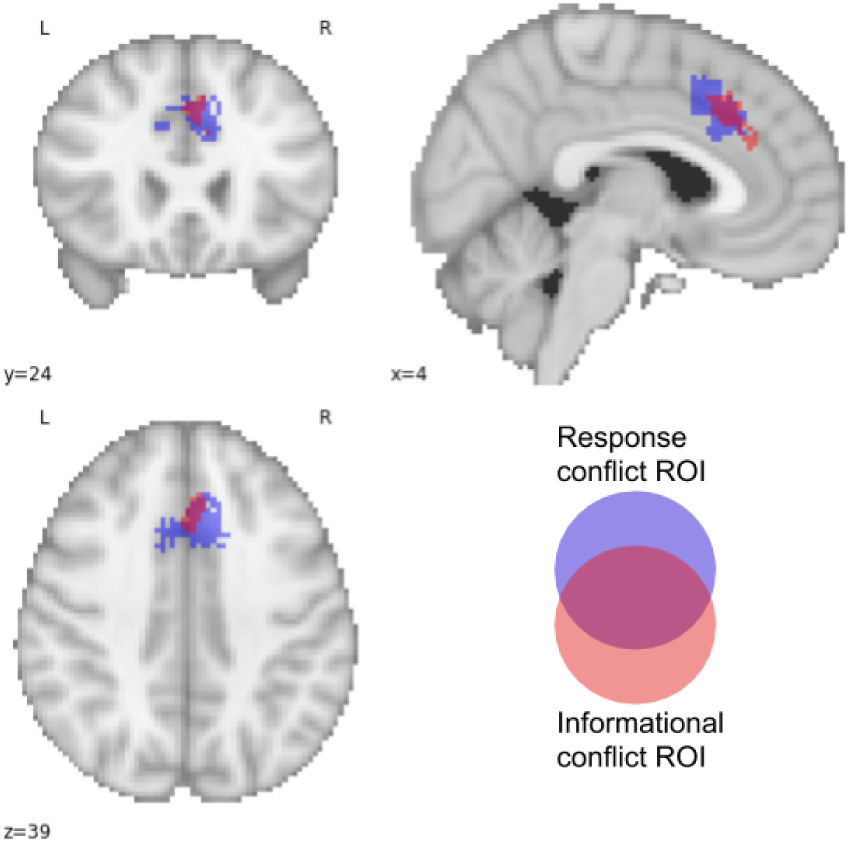
ROIs used for our confirmatory ROI analyses of the effect of religiosity on response conflict and informational conflict.

### Statistical Models

We applied hierarchical Bayesian models for all hypotheses to accommodate the hierarchical structure of the behavioral and fMRI data, with trials nested within participants. We constructed hierarchical Bayesian models with varying intercepts and varying slopes using the R package `brms` (Bürkner, 2017), which relies on the programming language `Stan` (Carpenter et al., 2017). This package incorporates `bridgesampling` (Gronau, Singmann, & Wagenmakers, 2017) for hypothesis testing by means of Bayes factors (BF) and posterior probabilities. The general form of our multilevel regression models is:

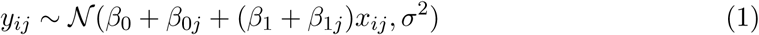

where *y*_*ij*_ is the outcome per trial per participant, and *x*_*ij*_ the corresponding value of the predictor. The subscript *i* is for the individual trials (*i* = 1…*n*_trials_) and the subscript *j* is for the participants (*j* = 1…*N*).

#### Prior Specification

We note that the most relevant parameter for making inferences in our specified models is the *β*_1_, i.e., the beta-weight for the (standardized) predictors of interest (e.g., Stroop condition, religiosity). As this parameter is used in the critical tests for our hypotheses, it is important to set appropriate priors particularly for this parameter. We chose *β*_1_ ∼ 𝒩 (0, 1) for the (standardized) predictors. This prior is listed as a recommended ‘generic weakly informative prior’ in the Stan manual (Betancourt, Vehtari, & Gelman, 2015), and has been used in this context before (e.g., Gelman, Lee, & Guo, 2015).

On the remaining parameters we used weakly-informative priors, whereby the priors for the regression weights (*β*’s) are derived from a normal distribution, and the priors on the scale parameters from a half-Cauchy distribution (𝒞^+^; Gelman, 2006): *β*_0_ ∼ 𝒩 (0, 10) for the fixed intercept; 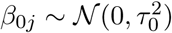 for the varying part of the intercept per participant; 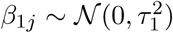 for the varying part of the predictor effect per participant; *τ* ∼ 𝒞^+^(0, 2) for the participant-level variance. Finally, we used the default LKJ-correlation prior to model the covariance matrices in hierarchical models (Lewandowski, Kurowicka, & Joe, 2009). That is, we used Ω_k_ ∼ LKJ(*ζ*), with Ω_k_ being the correlation matrix and *ζ* set to 1.

#### Interpretation of Evidence

Hypothesis testing was done by means of Bayes factors that evaluate the extent to which the data is likely under the alternative hypothesis (e.g., ℋ_1_–ℋ_8_) versus the corresponding null hypothesis ℋ_0_. The Bayes factor (BF) reflects the change from prior hypothesis or model probabilities to posterior hypothesis or model probabilities and as such quantifies the evidence that the data provide for ℋ_1_ versus ℋ_0_, reflected by:

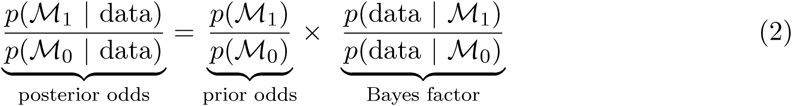

where ℳ_1_–ℳ_8_ and ℳ_0_ represent the models specified for ℋ_1_–ℋ_8_ and ℋ_0_, respectively. The Bayes factor BF_10_ then represents the ratio of the marginal likelihoods of the observed data under ℳ_1_ and ℳ_0_:

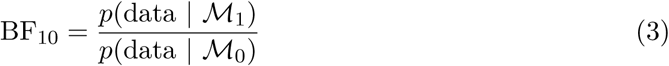

As our hypotheses are directed, we computed order-restricted Bayes factors, i.e., BF_+0_ in case of an expected positive effect. Note that the subscripts on Bayes factor to refer to the hypotheses being compared, with the first and second subscript referring to the one-sided hypothesis of interest and the null hypothesis, respectively. BF_+0_ is used in case of a hypothesized positive effect for the reference group or a positive relation between variables; BF_−0_ is used for a negative effect for the reference group or a negative relation between variables. As Bayes factors are fundamentally ratios that are transitive in nature, we can easily compute an order restricted Bayes factor; by (1) using the BF for the unrestricted model versus the null model, and (2) comparing the unrestricted model to an order restriction, we can then (3) use the resulting BFs to evaluate the order restriction versus the null model (Morey, 2015).

By default, prior model odds were assumed to be equal for both models that are compared against each other. As the evidence is quantified on a continuous scale, we also present the results as such. Nevertheless, we included a verbal summary of the results by means of the interpretation categories for Bayes factors proposed by Lee and Wagenmakers (2013, p. 105), based on the original labels specified by Jeffreys (1939). In addition to Bayes factors, we present the posterior model probabilities that are derived from the generated posterior samples.

For all outcome neutral tests we preregistered that a Bayes factor of at least 10 –the minimum value for strong evidence– was required to meet the criteria.

We declare that all models that are described below were constructed before the data were inspected. Additionally, all analyses were run as preregistered. Any deviations are explicitly mentioned in the manuscript.

## Results – Outcome Neutral Tests

### Behavioral Stroop Effect – Accuracy

A hierarchical logistic regression model with varying intercepts for the participants and a varying slope for the effect of Stroop congruency was constructed to model response accuracy. In order to validate the presence of a congruency effect on accuracy, i.e., a Stroop effect, we compared the model for ℋ_0_ containing only the varying intercept, to the model for ℋ− containing the varying intercept and the negative effect of congruency. ℋ− thus indicates that the incongruent condition *decreases* the probability of responding correctly on the Stroop task, relative to the congruent condition.

Results revealed a Bayes factor of 8.43 × 10^11^ in favor of the alternative model (ℳ_−_) relative to the null model (ℳ_0_). That is, BF_−0_ = 8.43 × 10^11^, indicating that the data are about 10^11^ times more likely under the model assuming lower accuracy for incongruent Stroop trials than for congruent Stroop trials. In order words, the data provide strong evidence for the Stroop effect indexed by accuracy (ℋ_1_). See Table 2 for a summary of the results of all four outcome neutral tests.

**Table 2.**
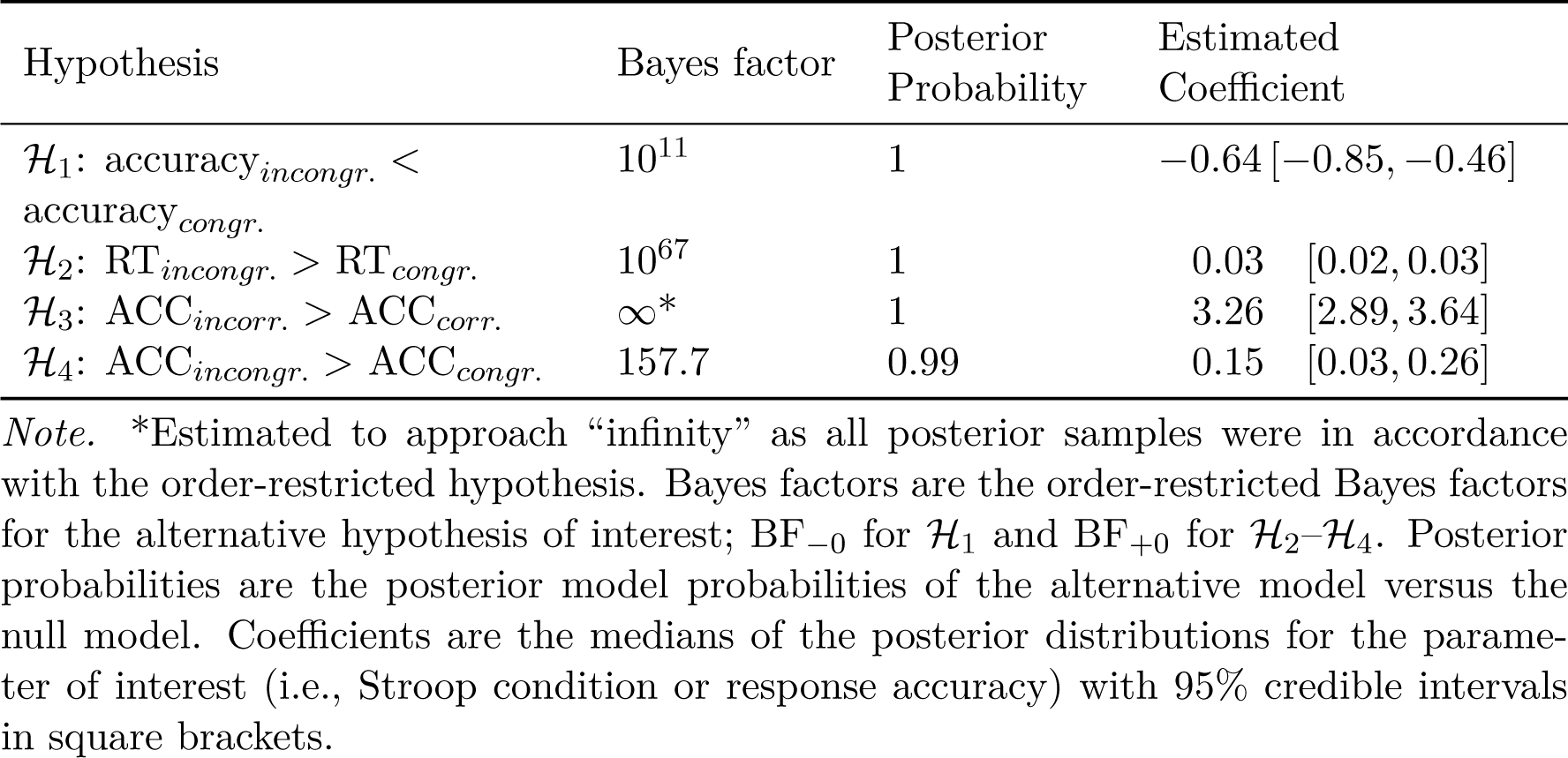
Results Outcome Neutral Tests.

### Behavioral Stroop Effect – Response Times

We used a similar hierarchical regression model with varying intercepts for the participants and a varying slope for the effect of Stroop condition to model reaction times. Note that only correct trials are included in the RT analysis. To account for the typical positive skew in RT data, we modelled reaction times as an ex-Gaussian distribution, i.e., a mixture of a Gaussian and an exponential distribution, which has been shown to fit empirical RT data well (Balota & Spieler, 1999; Balota & Yap, 2011; Whelan, 2008). This distribution is incorporated in the brms package, and thus only needed to be specified. Here we expected RTs to be longer for incongruent vs. congruent trials, hence the Bayes factor *BF*_+0_ was calculated for ratio between the marginal likelihoods of the observed data under ℋ_+_ versus ℋ_0_. Again, we expected a Bayes factor of at least 10.

We obtained a Bayes factor of 3.53 × 10^67^ in favor of ℳ_+_, that is BF_+0_ = 3.53 × 10^67^. In other words, we collected strong evidence for the Stroop interference effect on reaction times (ℋ_2_).

### Neural Processing – Response Conflict

The hierarchical nature of the fMRI data –being derived from multiple trials– was already taken into account in the calculation of the ‘incorrect–correct’ contrast and the ‘incongruent–congruent’ contrast in FSL: we exported the beta-values for each contrast per participant, as well as the variance for the contrasts, i.e., 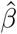 and 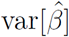. The inclusion of the variance parameter in the Bayesian models is important, because it allows one to retain the uncertainty associated with the activation level contrast, which is typically lost or ignored when extracting fMRI data for ROI-analyses.^10^ In order to test ℋ_3_ that the average contrast of ACC activation – the average ‘intercept’ or 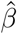 – was substantially different from 0, we used the function hypothesis which allows for directed hypothesis test of the specified parameters.^11^ 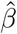 is calculated as 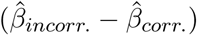, therefore the hypothesis states that 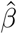 is larger than 0 (i.e., increased ACC activity for errors compared to correct responses). Here we calculated the Bayes factor for ℋ_+_ stating that 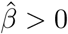.

We note that analyses that took the ‘incorrect–correct’ fMRI contrast as the dependent variable (ℋ_3_ and ℋ_7_) include data from 172 participants rather than 193, since some participants made no errors on the Stroop task.

The results showed evidence for the alternative hypothesis to approach “infinity”, that is BF_+0_ = ∞. Note that this Bayes factor was estimated by testing the proportion of posterior samples that satisfy the hypothesis that the intercept *>* 0. When all posterior samples are in accordance with the hypothesis, a Bayes factor of “infinity” can be obtained. In this case that means that the Bayes factor is at least 60, 000 since the model included 60, 000 samples. In other words, the neural data provide strong evidence that the ACC is sensitive to response accuracy on the Stroop task.

### Neural Processing – Informational Conflict

A similar procedure was used to test ℋ_4_, this time with the ACC activity contrast for Stroop congruency instead of response outcomes. That is, a hierarchical regression model with a varying intercept for the participants was constructed. The Bayes factor was calculated for the hypothesis that 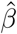 is larger than 0, since we expected 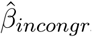. to be larger than 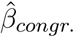, resulting in a positive contrast. Again, a Bayes factor of at least 10 was required to pass the outcome neutral criterion test.

A Bayes factor of 157.7 in favor of the alternative hypothesis was obtained (i.e., BF_+0_ = 157.7), indicating that the data provide strong evidence that the ACC is sensitive to informational conflict on the Stroop task.

The results of these four analyses indicate that all prespecified outcome neutral criteria were met.

## Results – Main Preregistered Analyses

### Behavioral Stroop Effect and Religiosity – Accuracy

In order to test ℋ_5_ whether self-reported religiosity of individuals is related to their performance on a conflict-inducing Stroop task, an extended Bayesian hierarchical logistic regression model was constructed, by adding religiosity as second-level predictor. Specifically, the model for ℋ_0_ included varying intercepts and varying slopes for Stroop condition (as before) per participant, plus the participant-level variables gender, age, and intelligence (i.e., the covariates). The model for the alternative hypothesis was identical plus the inclusion of religiosity as an additional participant-level predictor. Notably, an interaction term for religiosity × congruency was also included, as the effect of religiosity might be specific for performance in the conflict condition (i.e., the incongruent Stroop condition). As we expected a negative relation between religiosity and performance on the gender-Stroop task, we restricted the coefficient for religiosity to be negative in calculating the Bayes factor, i.e., we performed a one-sided test.^12^ The ratio of marginal likelihoods for the data under ℋ− versus ℋ_0_, i.e., the Bayes factor, were calculated to determine the evidence for the predictive value of religiosity in explaining Stroop performance.

A Bayes factor of 0.022 was obtained (i.e., BF_−0_ = 0.022, BF_0−_ = 44.8), indicating that the data provided more support for the null model than for the religiosity model. This result qualifies as strong evidence that religiosity is not negatively related to accuracy on the Stroop task. The posterior medians and the 95% credible interval for the coefficients of religiosity (−0.08 [−0.25, 0.09]) and of religiosity × Stroop condition (0.10 [−0.04, 0.24]) indicate that neither religiosity, nor the interaction between religiosity and Stroop condition was related to performance on the Stroop task (see also Figure 3a). The results of all main hypotheses are also summarized in Table 3. The parameters in the regression models for the four main analyses are displayed in the Appendix.

**Table 3.** Results Main Analyses.

| Hypothesis | Bayes factor | Posterior Probability | Estimated Coefficient |
| --- | --- | --- | --- |
| $\mathcal{H}_5$: Religiosity ↑ – Stroop performance (accuracy) ↓ | 0.022 (44.82) | 0.012 | $-0.08 [-0.25, 0.09]$ |
| $\mathcal{H}_6$: Religiosity ↑ – Stroop response times ↑ | $10^{-5}$ (25461) | 0.000 | $0.01 [-0.01, 0.02]$ |
| $\mathcal{H}_7$: Religiosity ↑ – ACC activity (response conflict) ↓ | 0.286 (3.49) | 0.172 | $-0.09 [-0.44, 0.26]$ |
| $\mathcal{H}_8$: Religiosity ↑ – ACC activity (informational conflict) ↓ | 0.046 (21.87) | 0.064 | $0.03 [-0.09, 0.15]$ |

*Note.* Bayes factors are the order-restricted Bayes factors for the alternative hypothesis of interest; $\text{BF}_{-0}$ for $\mathcal{H}_5$ , $\mathcal{H}_7$ , and $\mathcal{H}_8$ and $\text{BF}_{+0}$ for $\mathcal{H}_6$ . Evidence for the null hypothesis is given between brackets. Posterior probabilities are the posterior model probabilities of the alternative model versus the null model. Coefficients are the medians of the posterior distributions for the parameter of interest (i.e., religiosity) with 95% credible intervals in square brackets.

**Figure 3.**
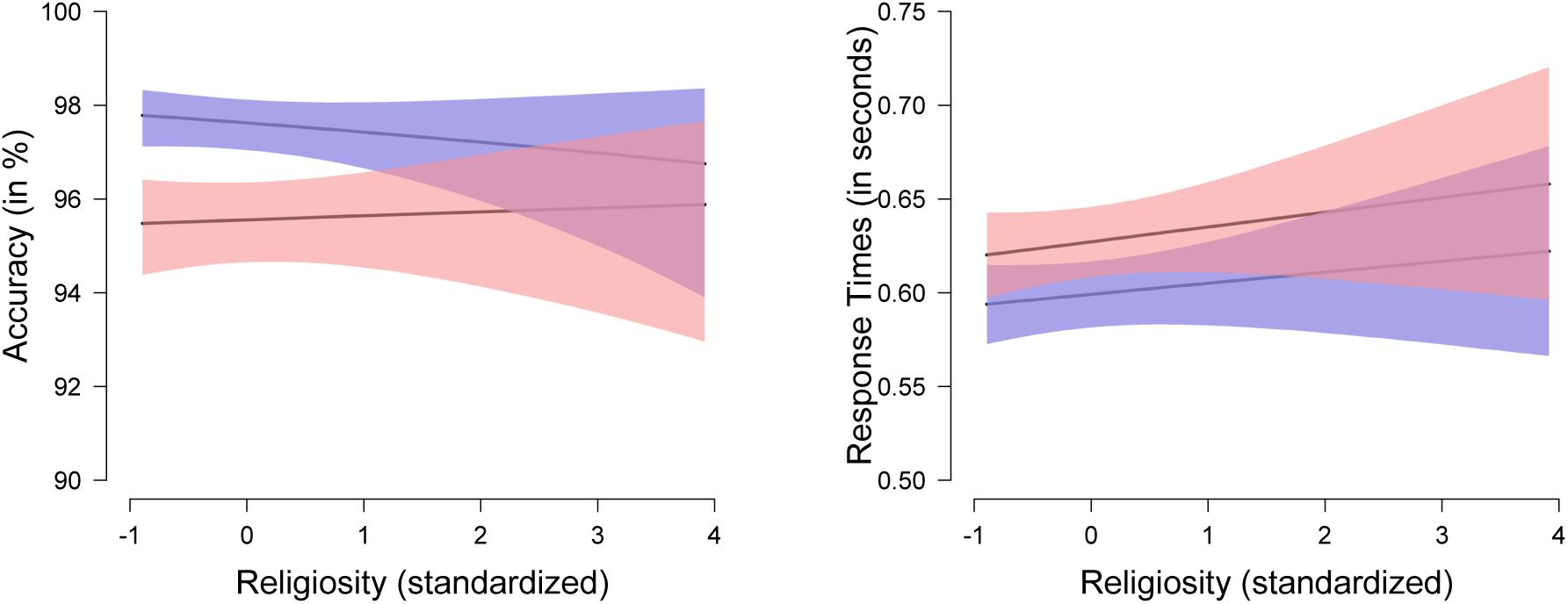
The marginal effect of religiosity on Stroop accuracy and response time, displayed per Stroop condition. The line with the blue 95% credible interval band indicates performance on congruent Stroop trials, the line with the red 95% credible interval band indicates performance on incongruent Stroop trials.

### Behavioral Stroop Effect and Religiosity – Response Times

We constructed a similar model with RT as the dependent variable; the model for ℋ_0_ was a hierarchical ex-Gaussian regression model for RT with varying intercepts and a varying slope for Stroop condition – including participant gender, age, and intelligence as covariates. For ℋ_+_, the model was identical with the added religiosity predictor and the religiosity × congruency interaction term. Again, we hypothesized that religiosity would be negatively related to Stroop performance, hence we expected a *positive* effect of religiosity on Stroop response times.

A Bayes factor of 3.93 × 10^−5^ was obtained (i.e., BF_+0_ = 3.93 × 10^−5^, BF_0+_ = 25461). Similar to the accuracy analysis, this indicates that the data do not provide support for the hypothesis that religiosity is related to longer response times on the Stroop task. Rather, we obtained strong evidence for the null hypothesis. The posterior medians for the coefficients of religiosity (0.01 [−0.01, 0.02]) and of religiosity × Stroop condition (0.00 [−0.00, 0.01]) corroborate that there was no main effect of religiosity on response times, nor was there an interaction of religiosity × Stroop condition on response times (see also Figure 3b).

### Neural Processing and Religiosity – Response Conflict

A Bayesian linear regression was performed in order to test ℋ_7_ whether self-reported religiosity is related to the ACC sensitivity to incorrect vs. correct responses on the Stroop task. The beta-values for the BOLD contrast in our specified ROI served as the dependent variable, i.e., the extracted 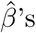. Again, the variance of the individual beta-values was included to take the uncertainty of the contrast estimation into account. Religiosity served as the predictor of interest and gender, age, and intelligence were added as covariates. That is, we compared the model including the contrast-intercept and the covariates (ℋ_0_) to the model additionally including the religiosity predictor. Based on the findings by Inzlicht et al. (2009), we expected a *negative* relation between religiosity and ACC activity induced by response conflict.

The results showed more evidence for the null model than for the model including religiosity as a predictor: BF_−0_ = 0.286 (i.e., BF_0−_ = 3.49). This Bayes factor is interpreted as moderate evidence against the hypothesis that religiosity is associated with reduced ACC sensitivity to response conflict in the Stroop task (i.e., the ‘incorrect–correct’ contrast). The posterior median and credible interval for the religiosity predictor were −0.09 [−0.44, 0.26]. The scatterplot in Figure 4a illustrates the (absence of an) association between religiosity and sensitivity of the ACC to response conflict.

**Figure 4.**
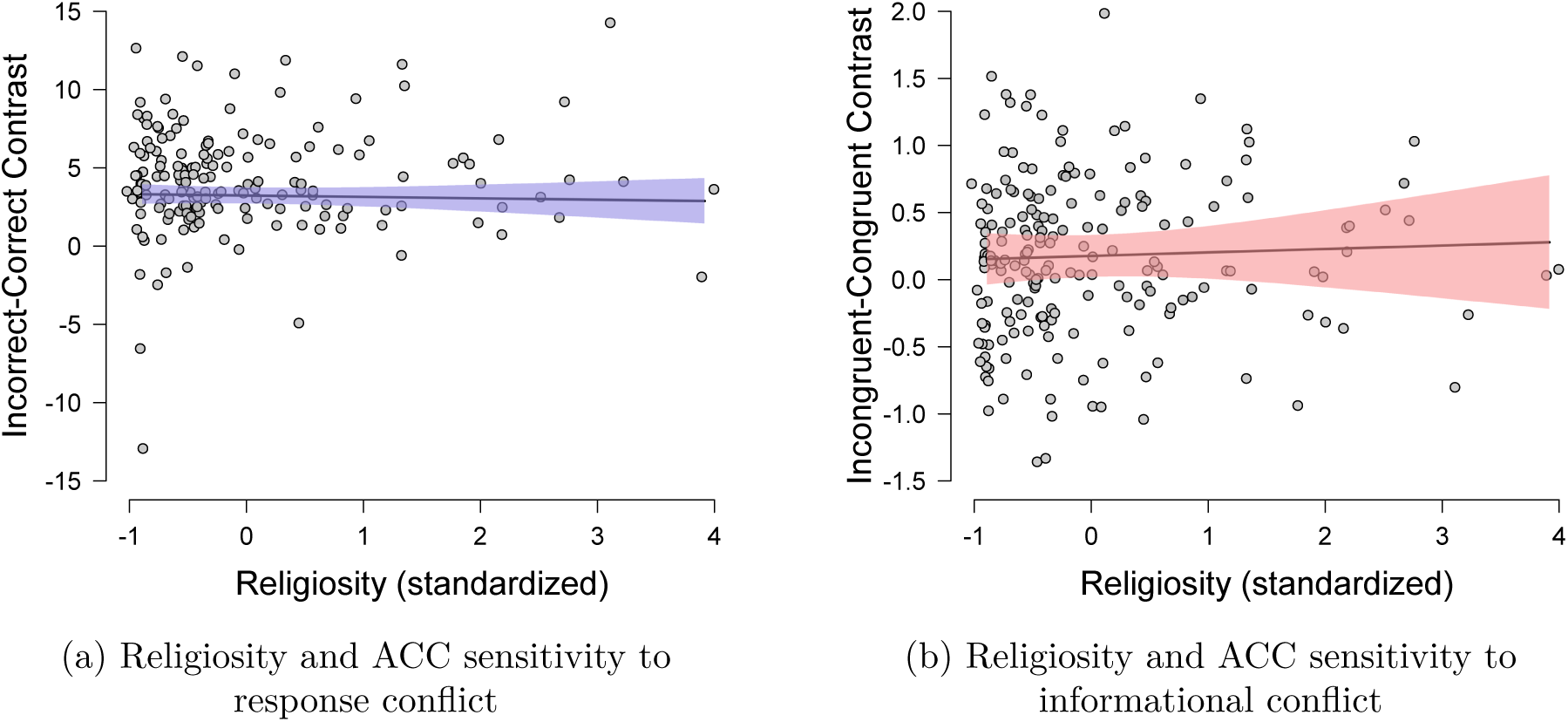
The relation between religiosity on the BOLD signal contrast for incorrect vs. correct responses on the Stroop task (left panel) and on the BOLD signal contrast for incongruent vs. congruent trials in the Stroop task (right panel). The plots display raw individual data points and Bayesian estimated linear effect of religiosity on the conflict-induced BOLD contrasts with 95% credible interval bands.

### Neural Processing and Religiosity – Informational Conflict

The same model comparison was performed with regard to the stimulus congruency contrast (i.e., ℋ_8_). Here, we used the 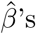’s of the incongruent–congruent BOLD contrast as the dependent variable. Again, we expected ACC activity to be negatively related to religiosity, while taking into account the effects of gender, age, and intelligence.

A Bayes factor of 0.046 (BF_−0_ = 0.046, BF_0−_ = 21.9) was obtained, indicating that the data provide strong evidence against the hypothesis that religiosity is related to reduced ACC sensitivity to informational conflict in the Stroop task (i.e., the ‘incongruent– congruent’ contrast). The posterior median and credible interval for the religiosity predictor were 0.03 [−0.09, 0.15]. The scatterplot in Figure 4b illustrates the (absence of an) association between religiosity and sensitivity of the ACC to informational conflict.

## Results – Exploratory Whole-Brain Analyses

In addition to the confirmatory ROI analyses, we conducted an exploratory (non-parametric) whole-brain analysis of the effect of religiosity on both response conflict and informational conflict. In addition, we ran an ‘intercept-only’ model (estimating the average effect of response and informational conflict) as an outcome neutral test. All whole-brain *t*-value maps and associated ‘1-*p*-value’ maps can be viewed at and downloaded from Neurovault (https://identifiers.org/neurovault.collection:6139).

### Outcome Neutral Tests

In Figure 5, we visualized the whole-brain results (as *t*-values) of the ‘intercept-only’ model for both the response conflict data (i.e., using the ‘incorrect–correct’ contrast; Figure 5A) and the informational conflict data (i.e., using the ‘incongruent–congruent’ contrast; Figure 5B).

**Figure 5.**
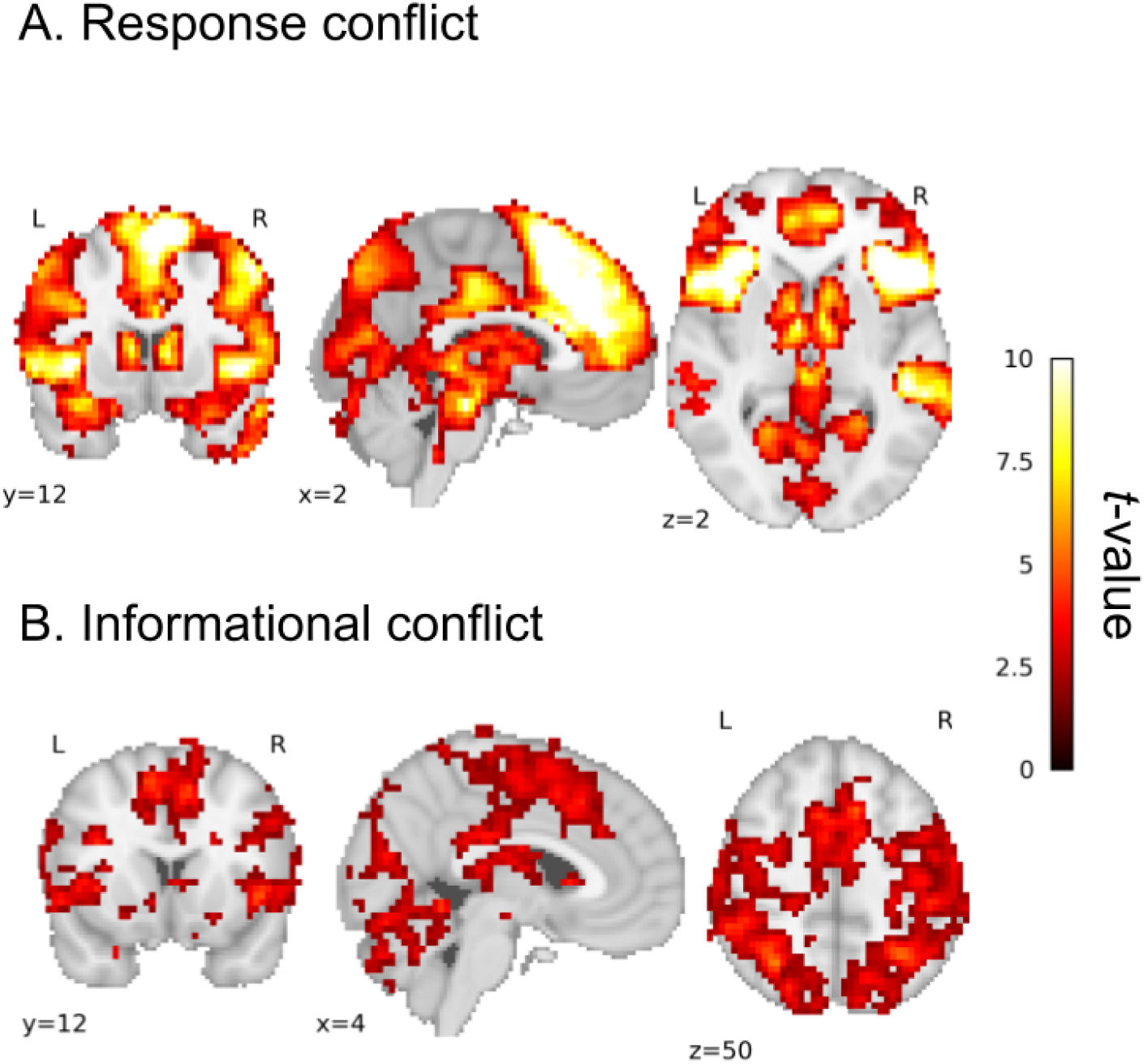
Brain maps with *t*-values corresponding to the outcome neutral (‘intercept-only’) test for both the (A) response conflict analysis and (B) informational conflict analysis. The brain maps were masked using *p*-values computed using FSL’s *randomize* with threshold-free cluster enhancement, which we thresholded at *p <* 0.05.

Both whole-brain maps show widespread effects in areas known to be involved in error monitoring and cognitive conflict (such as the ACC and insula). Note that the effects (i.e., *t*-values) are much larger in the response conflict analysis, presumably due to the relatively high variance in the first-level analysis stage due to high predictor correlation.

### Neural Processing and Religiosity – Response Conflict

After multiple comparison correction, no voxels were significantly associated with religiosity in the response conflict analysis.

### Neural Processing and Religiosity – Informational Conflict

Similar to the response conflict analysis, no voxels were significantly associated with religiosity after multiple comparison correction in the informational conflict analysis.

## Discussion

In the current preregistered study we investigated whether religiosity is associated with a reduced sensitivity to cognitive conflict as measured through behavioral performance on the Stroop task and neural activation in the anterior cingulate cortex (ACC). The data from the outcome neutral tests provided strong evidence that the gender-Stroop task induced cognitive conflict at the behavioral level (ℋ_1_ and ℋ_2_) and that this was reflected in increased ACC activity. The neuroimaging data showed that the ACC was responsive to both response conflict (incorrect vs. correct responses; ℋ_3_) and informational conflict (incongruent vs. congruent trials; ℋ_4_). However, individual differences in religiosity were not related to performance on the Stroop task as measured in accuracy (ℋ_5_) and response times (ℋ_6_). We also did not observe the hypothesized relation between religiosity and neural activation related to response conflict (ℋ_7_) or informational conflict (ℋ_8_). Overall, we obtained moderate to strong evidence in favor of the null hypotheses according to which religiosity is unrelated to sensitivity to cognitive conflict. Exploratory whole-brain analyses similarly showed that conflict-induced neural activity was not associated with religiosity.

These results cast doubt on the theoretical claim that religiosity is related to a reduced process of conflict sensitivity. Although this idea is central to various theories about religious beliefs (e.g., van Elk & Aleman, 2017; Inzlicht & Tullett, 2010; Schjoedt et al., 2013), our study shows that religious believers are not characterized by a *general tendency of* attenuated conflict sensitivity. An important motivation for conducting the current study was to address and overcome the limitations of previous studies in the field. We did so by increasing statistical power (i.e., we used a large sample) and by minimizing degrees of freedom (i.e., we preregistered all hypotheses, methods, and analyses and a priori specified a region of interest (ROI) for the fMRI analysis). Moreover, we curtailed the possibility of (unconscious) biases, as we separated the preprocessing of the fMRI data from the statistical analysis and only combined the fMRI data with the critical variable of interest (i.e., religiosity) in the final analysis steps.

Our null findings are perhaps not surprising in light of the recently voiced concerns about the replicability and reliability of neuroscientific findings, often related to problems of insufficiently powered studies (Button et al., 2013; Cremers, Wager, & Yarkoni, 2017; Szucs & Ioannidis, 2017) and general challenges in studying individual differences using neuroimaging (Dubois & Adolphs, 2016). For instance, Boekel et al. (2015) attempted to replicate 17 findings relating behavior to brain structures and found convincing evidence for only one out of the 17 included effects. Similarly, van Elk and Snoek (2019) recently failed to find support for the hypothesized relation between religiosity and grey matter volume in several brain areas that were identified in the literature as being associated with religiosity.

We also did not find behavioral evidence for impaired nor for enhanced Stroop performance among religious believers. This might reflect that religiosity is unrelated to low-level cognitive control processes. At the same time, the null finding may also reflect the paradox that highly robust experimental effects –such as the Stroop effect– are often difficult to relate to reliable individual differences, irrespective of the specific individual difference construct of interest (Hedge, Powell, & Sumner, 2018; Rouder, Kumar, & Haaf, 2019). That is, because these effects are very robust and automatic (“everybody Stroops”), the between-subjects variability is by definition relatively small. For correlational designs, this ‘problem’ of small between-subjects variability is further complicated by the presence of measurement error. Rouder et al. (2019) demonstrated that the ratio of true variability (i.e., true differences between individuals) to trial noise (i.e., measurement error) is 1 : 7. This unfavorable ratio renders the mission to uncover individual differences in cognitive tasks difficult, if not even impossible. Hierarchical models could mitigate these problems, as these models minimize the effect of trial noise by pulling the trial-level estimates toward the individual’s mean effect (known as hierarchical shrinkage). In the current study, we did apply hierarchical modeling for the response time models, as well as the neural ACC models (incorporated in the first-level fMRI models in FSL and by adding the variance parameter of the beta’s in the statistical models). Nevertheless, as acknowledged by Rouder et al. (2019), characterizing the degree of measurement error does not imply that the real underlying individual differences can be recovered. This casts doubt on the feasibility to detect true individual variation in cognitive control tasks, and hence to uncover associations with other measures. For example, Hedge et al. (2018) reported correlations of Stroop performance with other measures of cognitive control (e.g., Flanker task, Go/No-go task) ranging from −.14 to .14, none of which were significant. If we cannot even establish correlations between two tasks designed to measure exactly the *same* underlying phenomenon (i.e., cognitive control), the quest for reliable correlations between Stroop performance and more distant constructs such as religiosity seems all the more futile.

Although we obtained moderate to strong evidence for all null hypotheses related to religiosity and cognitive conflict, the current study does not imply that we should reject the notion of reduced conflict sensitivity as a defining characteristic of religious beliefs all together. It could well be that the relationship between religiosity and conflict sensitivity is restricted to specific instances or contexts and hinges strongly on the specific measures and operationalizations that are used. For example, in the study by Good et al. (2015) participants read a sermon about different qualities of God and then performed a Go/No-Go task with alcohol-related stimuli for which responses should be inhibited. As all participants refrained form alcohol consumption in their daily lives based on religious grounds, errors on the Go/No-Go task were seen as ‘religious’ errors, exposing participants’ ostensible pro-alcohol tendencies. The results showed that emphasizing the loving and forgiving nature of God reduced the ERN amplitude in response to religious errors, while emphasizing divine punishment did not affect the ERN compared to a control condition. In other words, it could well be that when participants first contemplate on the comforting nature of their religious beliefs, this may reduce conflict-related ACC activity as induced by a task that includes religion-relevant items and responses. Such a task has much higher ecological validity than the Stroop task that we employed in the current study following the work by Inzlicht et al. (2009). Similarly, the observed reduction of activity in religious believers’ DLPC and ACC while listening to a charismatic religious authority (Schjoedt & Bulbulia, 2011), may specifically depend on the religious content of the speech (and may disappear when the same religious authority would talk about public transport or gardening). It is thus important to do justice to the subjective nature of religious practices and experiences, when studying these topics. This resonates with concerns about the lack of ecological validity in many neuroscience studies on religion (e.g., Schjoedt & van Elk, in press): while studies such as the present one offer high experimental control, the measures do not capture the ‘true stuff’ that most psychologists and neuroscientists of religion are interested in, namely lived religious beliefs and experiences.

We see two broad future directions for the field. First, the development of new and sophisticated techniques in neuroscience could allow for interesting new hypotheses and measures. For instance, the use of multi-voxel pattern analysis (MVPA) may provide insight into the representational nature of religious concepts endorsed by believers; a question could be whether the neural representations of religious agents such as ‘God’, ‘angels’, or ‘Satan’ are more similar to real people such as ‘Napoleon’ and ‘Donald Trump’ or to imaginary agents such as ‘Santa Claus’ and ‘Superman’ (cf. Leshinskaya, Contreras, Caramazza, & Mitchell, 2017).

Novel methods for assessing brain connectivity also allow for the investigation of new questions. (e.g., Huntenburg, Bazin, & Margulies, 2018; Margulies et al., 2016) One could assess for instance the relationship between religiosity and the integration of information from sensory cortical areas and the default mode network (DMN), a network that is implicated in abstract, high-level thinking. A hypothesis could be that religious believers are more likely to show a dissociation between the DMN and primary sensory areas. This could be studied in a correlational resting-state design, or alternatively, one could assess believers’ brain connectivity while engaging in contemplation of their (religious) beliefs or actions. For instance, intense personal prayer may be associated with a decoupling of internal self-referential processing in the DMN and perceptual processing in the sensory cortices specifically during the prayer experience, similar to what was found for shamanic trance-experiences (Hove et al., 2015).

Second, and relatedly, we believe there is much promise in future endeavours that focus on the application of paradigms and tasks that have higher ecological validity and more closely implicate religious concepts, as in the examples given above. Such an approach can hopefully do more justice to the multifaceted nature of religious beliefs and practices and can pave the way for a truly better understanding of the mechanisms and processes involved in religiosity.

## Appendix A

### Population Imaging of Psychology project

The data for this study was collected as part of the Population Imaging of Psychology project (PIoP), which was conducted at the Spinoza Center for Neuroimaging at the University of Amsterdam. The aim of the PIoP was to offer researchers the opportunity to collect brain-imaging data from a large sample of participants (intended N = 250), in association with their individual difference measure of interest. The data was collected between May 2015 and April 2016.

Standard measurements that were collected for each participant included a structural T1 MRI scan, task-free resting state fMRI (6 minutes), a diffusion tensor imaging (DTI) scan, and different functional localizer scans that were collected using EPI sequences, including the Gender Stroop task, an emotional matching task (Hariri, Bookheimer, & Mazziotta, 2000), a working memory task (Pessoa, Gutierrez, Bandettini, & Ungerleider, 2002), and the anticipation of positively vs. negatively valenced stimuli (Oosterwijk, 2017). In addition, demographic variables were recorded (gender, age, socio-economic status) for each participant, as well as two personality questionnaires, the NEO-FFI (Costa & McCrae, 1992) and the SCID (First, Gibbon, Spitzer, & Benjamin, 1997), and an intelligence test (Raven’s matrices; Raven, 2000). Finally, measures on religiosity and religious experiences were included (see Methods for details on the religiosity scale that was used in the present study).

#### Additional Religiosity Items

1. To what extent do you consider yourself to be spiritual?
2. To what extent do you believe in paranormal phenomena (e.g., astrology or telepathy)?
3. To what extent are your parents religious?
4. To what extent do your parents frequently visit a church or religious meeting?
5. Do your parents have a religious lifestyle (e.g., don’t go shopping on Sunday, pray before dinner)?

## Appendix B

### Coefficient Plots

**Figure B1.**
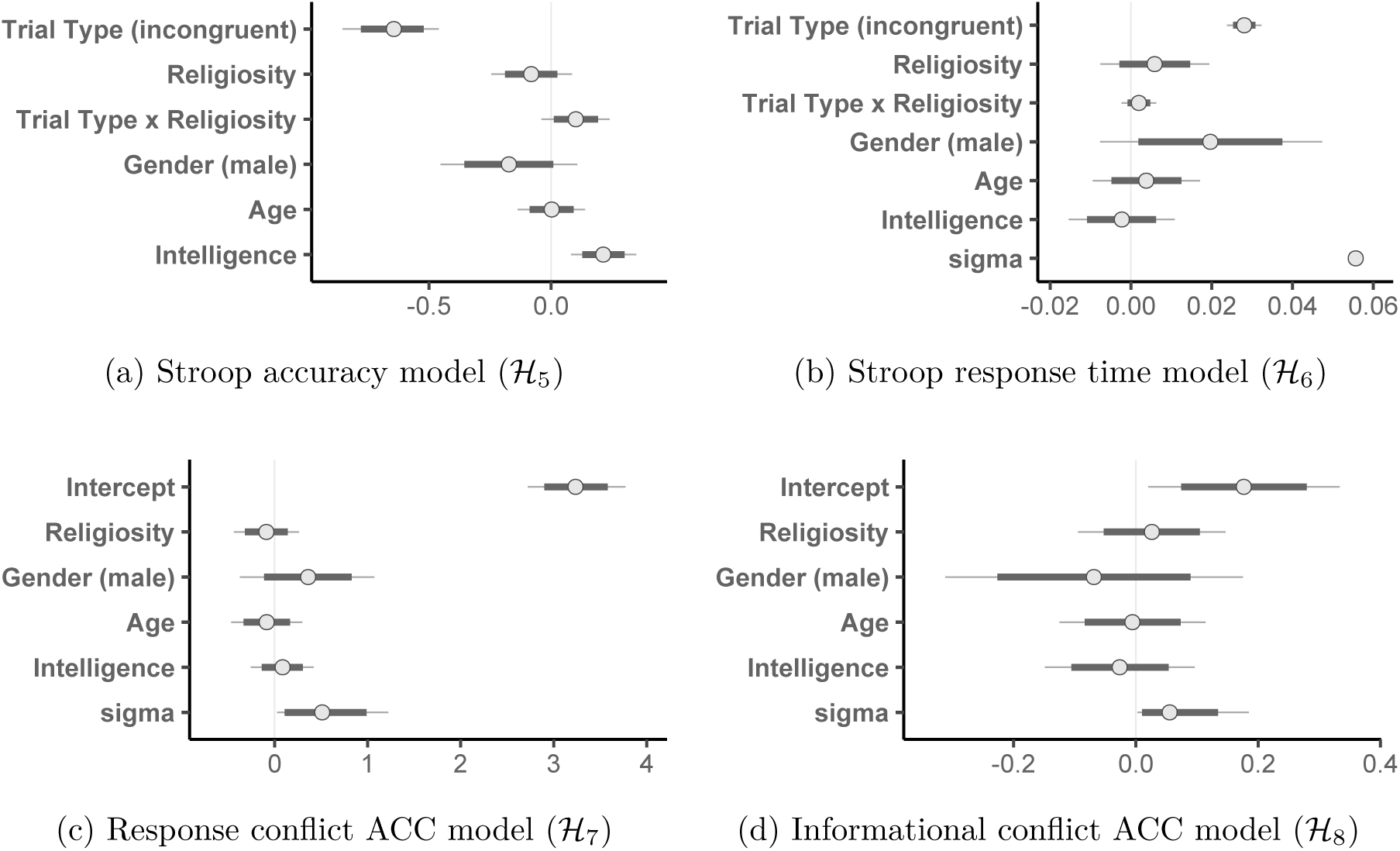
Coefficients of the fixed effects on Stroop accuracy (top left panel), Stroop response times (top right panel), response conflict ACC activity (bottom left panel), and informational conflict ACC activity (bottom right panel), derived from the Bayesian regression models. For each predictor, points represent the median estimates, thick lines the 80% credible interval and thin lines the 95% credible interval. Note that predictors in the accuracy model are on a linear scale and should be transformed by the inverse logit link to reflect probabilities. In the accuracy and response time models, the intercepts are omitted to enhance visibility of the predictors.

## Footnotes

1 Response conflict is here defined as the conflict between the actual and the correct response, rather than the prepotent and the correct response.

2 Based on the aforementioned theories addressing believers’ failure to notice incompatibility between different sources of contradicting information, we would primarily expect a negative association between religiosity and informational conflict (rather than response conflict). However, from an empirical perspective, our study most closely resembles the design by Inzlicht et al. (2009), who measured and obtained support for a relation between religiosity and neural markers of response conflict.

3 Specifically, LS was involved in data collection and (pre)processing the MRI data and has no access to the religiosity data. MvE and SH formulated the research questions and hypotheses without any access to the MRI data.

4 Of the 21 excluded participants, 19 made no errors and 2 participants made 1 error, but no reliable signal could be extracted for this error trial.

5 This meta-analysis reports a median sample size of approximately 22 for fMRI studies.

6 The face Stroop task - instead of the regular word-color variant - was chosen because it offers optimal opportunities for dissociating between perceptual processing of target and distractor dimensions, as processing of the distractor faces can straightforwardly be linked to activation patterns in the fusiform face area (FFA; Egner & Hirsch, 2005). In the current study, however, we were mainly interested in the cognitive conflict aspect rather than perceptual processing, and therefore solely focused on activation in the ACC.

7 The Dutch labels were ‘man’, ‘heer’,’vrouw’, and ‘dame’, respectively.

8 We note that the current design was suboptimal in estimating the effect of informational conflict (but not response conflict) in the fMRI data. Due to insufficient ‘jittering’ of the interstimulus interval, the first-level predictors for congruent and incongruent trails were strongly negatively corrected 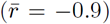. While this does not bias our results (the generalized least squares estimator we used is still unbiased), it *does* increase the variance of our first-level results, which in turn reduces the power of finding a correlation of religiosity with the first-level effect of informational conflict (operationalized by the ‘incongruent-congruent’ contrast). This issue only applies to the ‘incongruent-congruent’ contrast, not the ‘incorrect-correct’ contrast (as these predictors are much less correlated with each other, 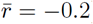).

9 These maps were generated on February 26th, 2019.

10 The possibility to include the variance of the observations in the regression model formula was added for the purpose of meta-analyses (Vuorre, 2016). However, it also serves the current purpose very well.

11 The term intercept may be somewhat confusing here. Since the outcome variable is the contrast between the incongruent and congruent condition (i.e., the difference), we only include the intercept in this model, and hence look at the effect of the parameter ‘intercept’.

12 The coefficient for the interaction term was not order-restricted.

